# True-atomic-resolution insights into the interactions of antibiotic rifampicin and rifamycin derivatives with orphan CYP143 of *M.tuberculosis*

**DOI:** 10.64898/2025.12.22.695974

**Authors:** Andrei Gilep, Christopher MacCarthy, Tatsiana Varaksa, Marianna Karpova, Irina Grabovec, Veronika Karpushenkova, Ivan Kapranov, Tatyana Sushko, Alena Karputs, Kouhei Tsumoto, Valentin Borshchevskiy, Natallia Strushkevich

## Abstract

Analysis of drug-resistant strains of *M.tuberculosis* revealed a mutation in cytochrome P450 CYP143 gene located in the ESX-5 secretion cluster. The function of CYP143 is unknown. Available synteny information is insufficient to straightforwardly infer potential CYP143 substrates, because a full set of effector proteins/substrates exported by the ESX-5 secretion system has not yet been identified. Here we found that CYP143 is highly conserved among mycobacteria. We leverage the possible association of the G334A mutation in CYP143 with the development of resistance to test the frontline antitubercular drug rifampicin using purified proteins. Binding of rifampicin does not induce typical P450 spectral response, while the crystal structure at atomic resolution reveals the binding mode in the active site. The drug binds above the heme and adopts “closed” conformation of the ansa chain running almost parallel to the naphthoquinone core. In this conformation the oxygen atom bridging C12 and C29 of the macrocycle makes a hydrogen bond with a water molecule coordinating heme iron. The same binding mode was observed in crystal structures in complex with rifaximin, rifamycin S and two synthetic derivatives. In the ligand-bound state the protein retains an open conformation even in the presence of the redox partner, as evident from the crystal structure of the ternary complex. Despite binding close to the heme, no conversion was observed showing that either the rifamycins or the reduction system, or both, are insufficient to support a full catalytic cycle. These results pave the way to understand the development of rifampicin resistance/tolerance and the role of orphan CYPome of *M.tuberculosis*.

## Introduction

The state of the global tuberculosis (TB) epidemic in 2023 revealed 10.8 million new cases. Drug-resistant TB remains a major challenge, with a significant gap in diagnosis and treatment for multidrug- and rifampicin-resistant TB (MDR/RR-TB), with only 44% diagnosed and treated (WHO Global Tuberculosis Report 2024). *Mycobacterium tuberculosis* (Mtb) has developed a wide array of mechanisms that impact tolerance and resistance to anti-TB drugs^1^. Understanding these mechanisms and identification of specific target/s is fundamental to the development of new therapeutic alternatives.

One of the mechanisms of drug resistance is drug metabolism by respective enzymes rendering the drug inactive. Unlike humans having Phase I (predominately cytochrome P450, CYP) and Phase II drug-metabolising enzymes, bacteria typically use different enzymatic strategies developed for antibiotic inactivation^2^, including hydrolysis, group transfer, and redox mechanisms. What is striking about the *M. tuberculosis* genome is its exceptionally high ‘density of CYP genes’ – approximately 240-fold greater than in humans. While the human genome (∼3000 Mb) encodes 57 CYPs, Mtb harbors 20 CYPs within only 4.41 Mb^3^, suggesting that many of these isoforms may play crucial and specialized roles in bacterial physiology. Actinomycetes to which Mtb belongs have numerous CYP genes known to produce natural compounds^4^. However, CYPome is dramatically reduced in obligate pathogens, for example, *Mycobacterium leprae* has only one CYP gene. The question of the function of the majority CYPs in Mtb, and whether they are involved in drug resistance remains open.

Whole genome sequencing of Mtb clinical isolates reveals single nucleotide variations (SNV) in CYPs that might affect drug susceptibility^5^. The G334A mutation in CYP143 is remarkably enriched in drug-resistant isolates compared to sensitive isolates, with a highly significant p-value of 0.000125^5^. This mutation appeared in 91% (10 out of 11) rifampicin mono-resistant, in 73% (52 out of 71) MDR, in 79% (23 out of 29) isoniazid mono-resistant, in 75% (6 out of 8) pre-XDR Mtb isolates^5^.

There is little information available about CYP143. The encoding gene *Rv1785c* is non-essential for *in vitro* growth of Mtb H37Rv^6,7,8,9.^ *Rv1785c* is not significantly induced or repressed according to transcriptomics data^10^, but its expression is elevated under rifampicin or isoniazid treatment^11, 12^. *In vivo* data about CYP143 as well as the majority of Mtb CYPs is not available. The essentiality of individual CYP genes can be complex and sometimes context-dependent. For example, while some CYP inhibitors show potent anti-mycobacterial effects, genetic knockout studies might show varied results depending on the specific growth conditions^3^. Some conditions are hard to model, like the dynamic lung microbiome, because it is a complex, highly individual system with constant microbial immigration and clearance^13^.

The CYP143 gene along with ferredoxin *Rv1786* are found within the ESX-5 cluster. *Mycobacterium sp*. has five specialized ESX export systems, ESX-1 to ESX-5 (also known as the Type VII secretion system) that are critical for growth and pathogenesis^14^. The ESX-5 is the most recently evolved cluster and has been linked to the drug resistance^15,16^. The genes encoding CYP143 and ferredoxin apparently do not belong to the secretion system, and if they were acquired by the slow-growing Mtb via plasmid mediated horizontal transfer as suggested for the ESX-5 gene cluster^17^,their beneficial traits might be synergistic with this particular secretion system. However, the function of CYP143 remains unclear.

In this study, we focused on identifying specific ligands for wild type and G334A mutant form of CYP143 and determining their binding mode using X-ray crystallography. Next, we solved the structure of the ternary complex CYP143-ferredoxin-ligand to determine whether the redox partner binding changes and/or enhances ligand binding. This is the first reported structure of the complex of the ligand–bound CYP with [Fe3S4]-type ferredoxin. Our data shed light on the fate of rifampicin and other related ansamycins within Mtb.

## Results

Phylogenetic analysis of mycobacterial CYPome revealed that CYP143 is closely related to CYP188 yet another orphan protein (Fig.1, A), which appears to have diverged from a common ancestor. The proteins with identified substrates (e.g., CYP124 and CYP125) are grouped in different branches highlighting the different evolution and specificity of CYP143 (Fig.1, B). CYP143 is conserved in the Mycobacterium tuberculosis complex (MTC) group, whereas CYP188 can be found in some pathogenic and nonpathogenic mycobacteria (Fig. 1C). We previously observed a strong coevolution within Mycobacteria for ferredoxin Rv1786 adjacent to CYP143^18^ (Fig. S1), suggesting a functional redox partner association. The pair is found in all slow-growing pathogenic Mycobacteria (Fig. 1C) and within a specific gene cluster encoding components of the ESX-5 secretion system (Fig. S2).

**Figure 1.**
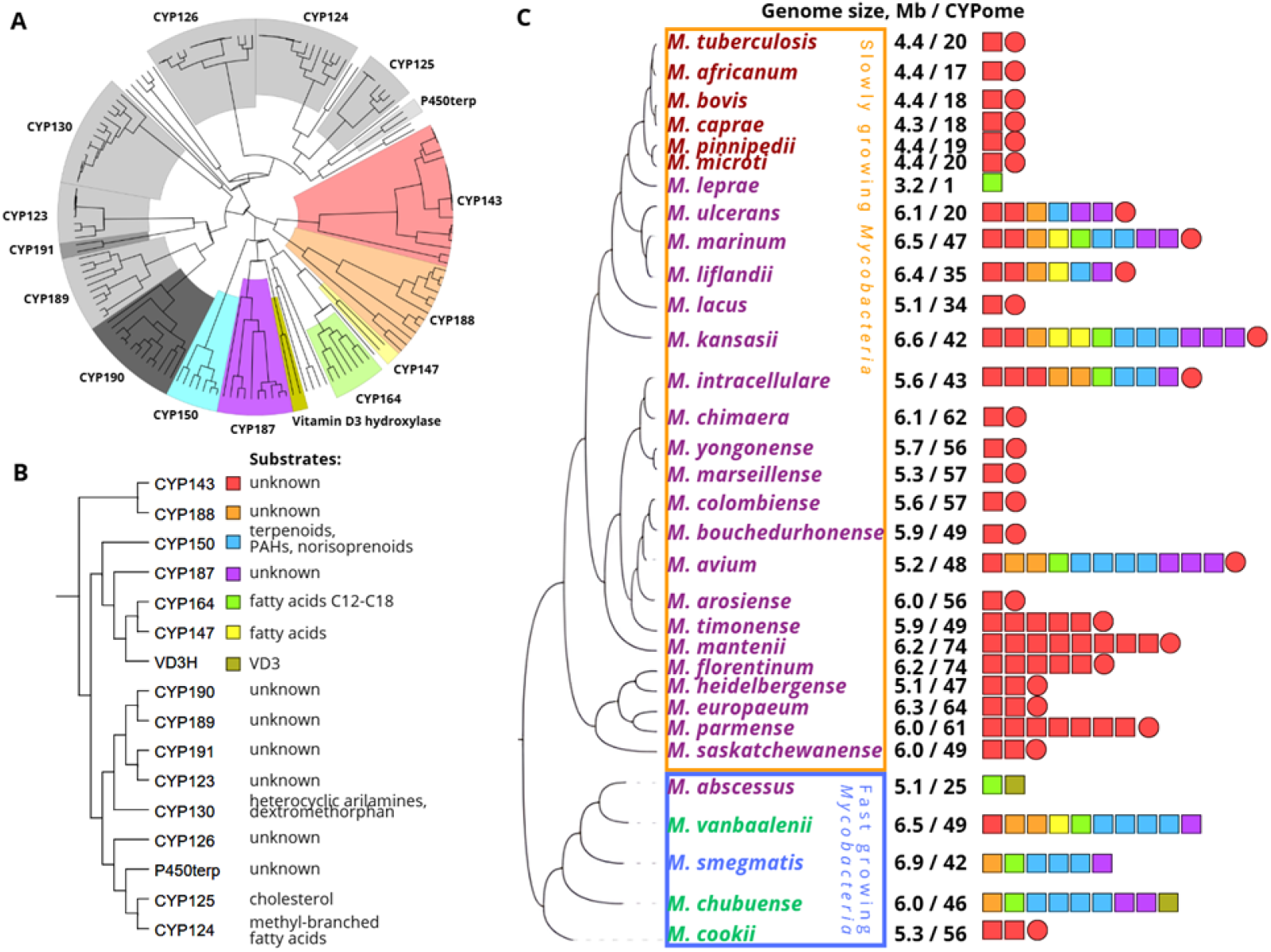
A ̶ Phylogenetic tree of P450 protein sequences from *Mycobacteria* (detailed in Figure S1 (in Supplementary)). B ̶ Phylogenetic relationships of CYPs. C ̶ Distribution of CYPs and Rv1786 across *Mycobacteria*. The colored squares depict the number of annotated in databases CYP isoforms, the filled circles - annotated ferredoxin Rv1786. Free-living nonpathogenic species highlighted in green, commensal species - in blue, species that cause diseases other than tuberculosis - in purple, and members of the MTC group - in red. Slowly growing *Mycobacteria* are separated from fast growing species by colored boxes. The number of CYP genes is provided for a given genome.

Surprisingly, CYP143 gene expression is not coregulated with the expression of ESX-5 genes under overexpression of different transcription factors^10^. Notably, in some species more than one CYP143 isoform is present (Fig. 1C), but in slowly growing bacteria only one copy is located within the ESX-5 gene cluster along with respective ferredoxin. In *M. leprae* CYP143 and Rv1786 are absent, but the ESX-5 system is only partially conserved^19,17^.

A variant search for CYP143 gene (*Rv1785c*) using genomic data of Mtb deposited to TB-portal (https://tbportals.niaid.nih.gov/depot) revealed 30 missense and 13 synonymous variants. The G334A missense variant seems to be quite common in different types of drug resistant isolates (Table 1). This data suggests a possible contribution of the G334A variant to the development of drug tolerance or resistance. It is also important to note that Gly residue in this position is conserved among slowly growing *Mycobacteria* (Fig. S3).

**Table 1.** The frequency of the CYP143 Gly334Ala single nucleotide variation (SNV) according to the genomic sequence data available in TB-portal.

| Drug resistance type | Number of isolates | SNV frequency |
| --- | --- | --- |
| HR-TB | 367 | 0.048404 |
| MDR-TB | 3417 | 0.450673 |
| Other | 342 | 0.045107 |
| Pre-XDR-TB | 2026 | 0.267212 |
| RR-TB | 60 | 0.007913 |
| Sensitive | 1223 | 0.161303 |
| XDR-TB | 147 | 0.019388 |

We synthesized the G334A mutant, expressed it in *E.coli*, purified to homogeneity and used it in our functional assays and structural studies. Qwing to strong absorbance of rifampicin (RIF) and its derivatives as well as unconventional mode of binding (see below) a spectrophotometric titration to determine Kd values was not applicable. The SPR using both His-tag and biotinylated proteins as well as ITC (Fig. S4), were also not successful for quantifying the interaction between rifamycins and CYP143. The SPR measures mass accumulation at the sensor surface upon the interactions between the immobilized protein and analyte, while the ITC relies on released or absorbed heat upon binding. In case of CYP143, immobilization likely affects protein flexibility and ultimately hinders the binding of ligands. As binding of RIF does not induce conformational changes (described below), ITC could not be successfully applied because of small enthalpy changes. Limited solubility of RIF and especially its derivatives in buffer solutions further complicates both methods.

Direct titration of rifampicin with CYP143 was carried out using a membrane-based separation step to remove unbound ligand, followed by centrifugation to isolate the protein–ligand complex. In the concentration range 15-60 μM of the CYP143 protein, we observed equimolar binding of RIF (initial concentration 100 μM) for both WT and mutant forms. No binding was observed for other Mtb CYPs – CYP51 and CYP141 (used as negative controls) which indicate specific interaction of CYP143 with RIF.

Of note, the seemingly low affinity of RIF for CYP143 may reflect a broader phenomenon in which certain ligands exhibit lower affinities for Mtb proteins than for their human counterparts. This difference can be influenced by a pronounced “concentration effect” that arises as molecules diffuse from the host macrophage to the intracellular, macrophage-residing bacilli. For instance, a single ligand molecule in a human alveolar macrophage corresponds to a concentration of ∼13.0 pM (1.3×10-11 M), given the macrophage volume of ∼4.99 pL (∼4.99×10-12L)^20^. In contrast, one molecule per Mtb cell corresponds to ∼0.24 μM (2.4×10-7 M), based on an average cell volume of ∼2.9 fL (∼2.9×10-16 L)^21^. Thus, even very small absolute molecule numbers translate into much higher effective concentrations at the scale of a single Mtb cell.

Azole compounds are promising antitubercular (anti-TB) agents, with some antifungal azoles demonstrating antibacterial activity against Mtb^22,23^. They are known binders to CYP proteins and typically induce type II spectral response by coordinating heme iron via nitrogen with lone pairs of electrons. We compared azole binding properties of the G334A mutant and wild type CYP143 (Table 2). The differences in binding are significant only for fluconazole and prochloraz highlighting the effect of mutation. Low-affinity enthalpy-driven binding was also detected using ITC for WT (Fig. S5 and Table S1). Overall, the binding affinity for commonly used antifungals is rather low for both mutant and WT proteins. Notably, hot spot mutations in *Candida albicans* that cause antifungal resistance occur in CYP51 (ERG11), in the same structural region. Well documented Gly→X substitutions in the heme-proximal loop (e.g., *C. albicans* G464S; *C. neoformans* G484S) reduce fluconazole affinity and confer azole resistance^24,25^.

**Table 2.**
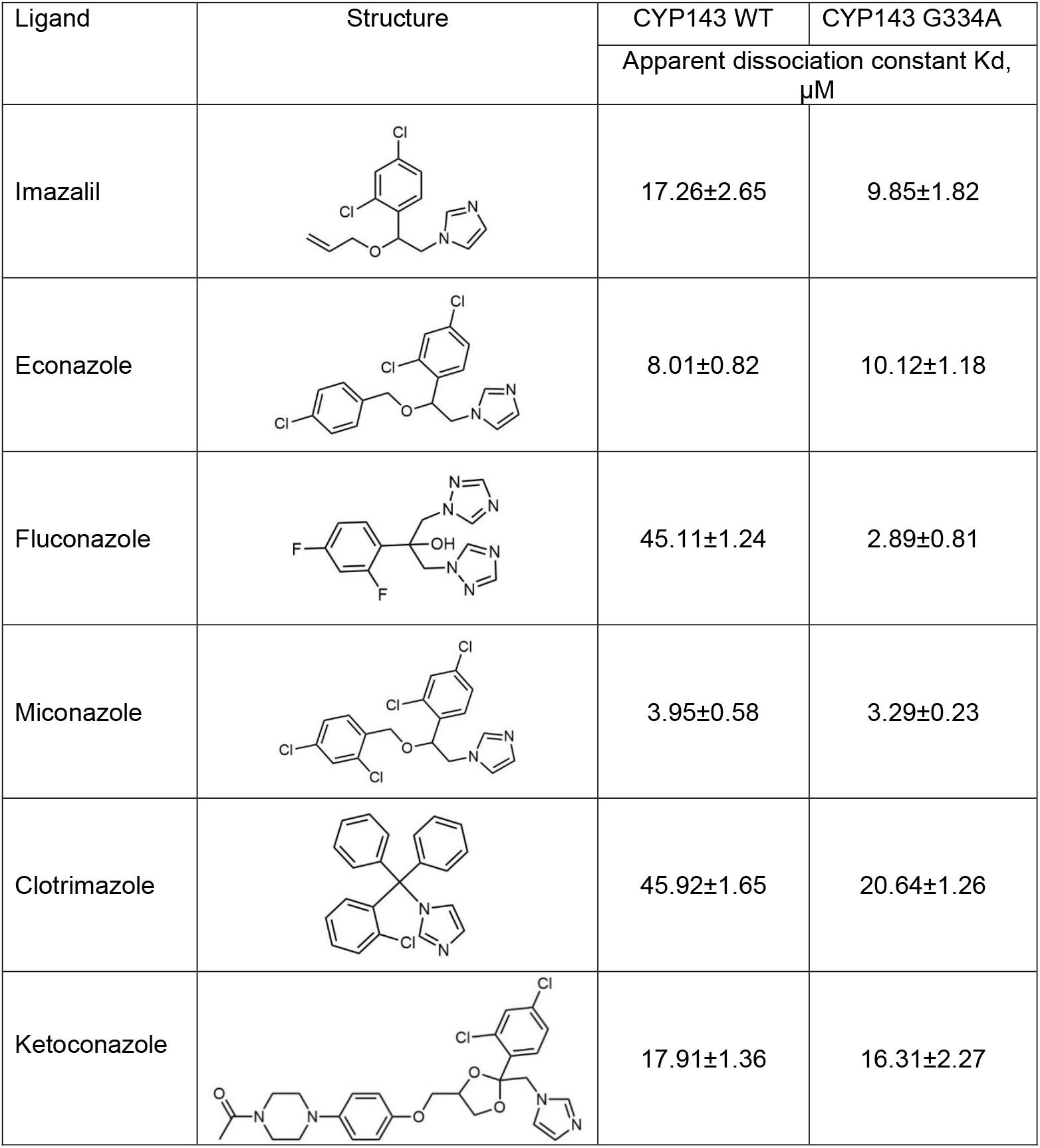
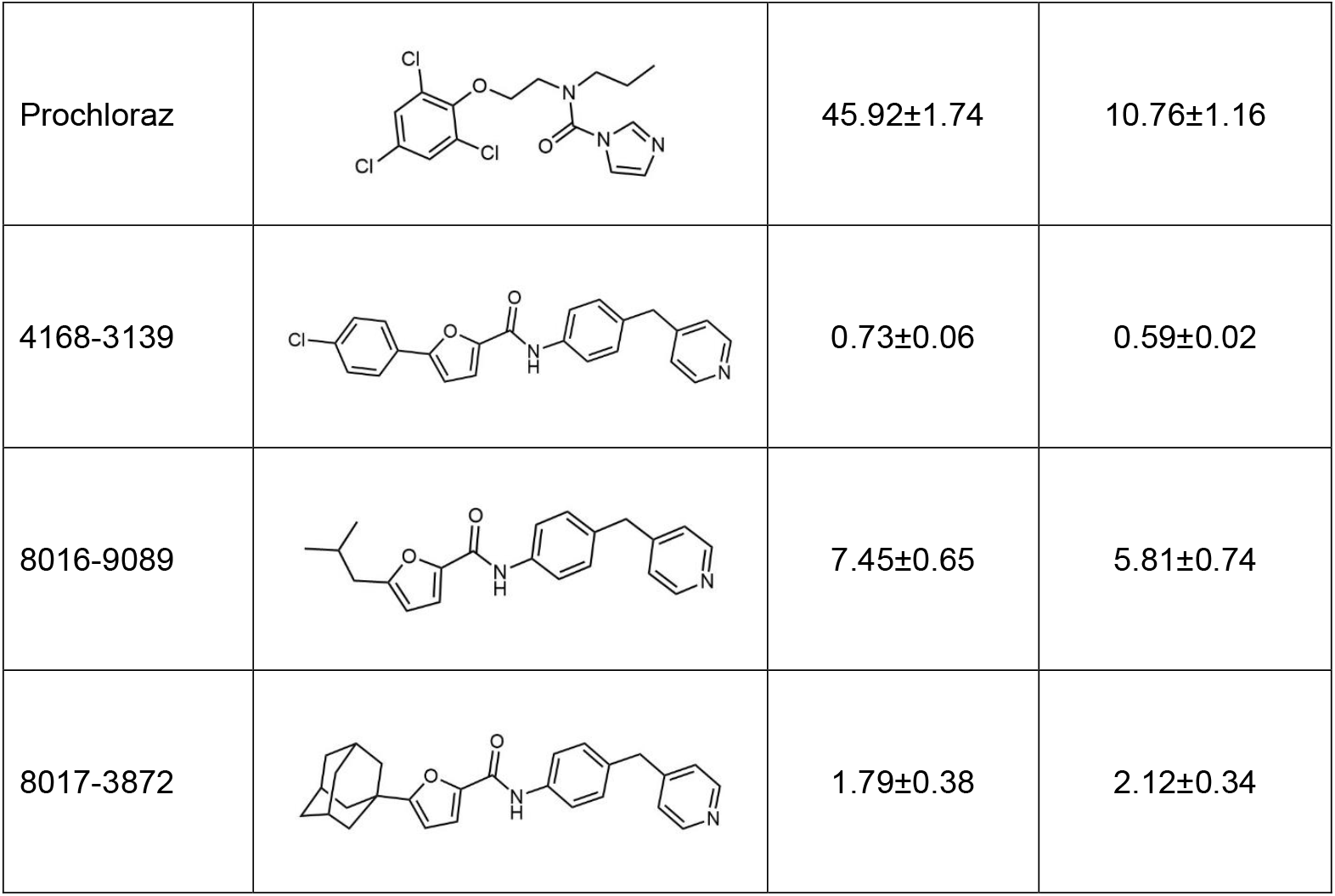
The results of spectral titration of CYP143 with different ligands.

| Ligand | Structure | CYP143 WT | CYP143 G334A |
| --- | --- | --- | --- |
| | | Apparent dissociation constant K<sub>d</sub>, μM | |
| Imazalil |  | 17.26±2.65 | 9.85±1.82 |
| Econazole |  | 8.01±0.82 | 10.12±1.18 |
| Fluconazole |  | 45.11±1.24 | 2.89±0.81 |
| Miconazole |  | 3.95±0.58 | 3.29±0.23 |
| Clotrimazole |  | 45.92±1.65 | 20.64±1.26 |
| Ketoconazole |  | 17.91±1.36 | 16.31±2.27 |
| Prochloraz |  | 45.92±1.74 | 10.76±1.16 |
| 4168-3139 |  | 0.73±0.06 | 0.59±0.02 |
| 8016-9089 |  | 7.45±0.65 | 5.81±0.74 |
| 8017-3872 |  | 1.79±0.38 | 2.12±0.34 |

We also tested several hits identified from HTS performed using WT protein and the library of 3800 compounds. Three compounds containing pyridine as a heme–iron binding moiety show higher affinity (Table 2) and the scaffold is similar to that identified in fragment-based screening^26^. Attempts to obtain crystal structure in complex with any type II ligand so far have been unsuccessful.

We reasoned that if the mutation appeared often in the rifampicin (RIF) resistant isolates, then this CYP might be involved in its binding and/or degradation. To test this hypothesis, we probed CYP143 for monooxygenase, demethylase, peroxidase and cleavage activity with RIF and rifamycin S as substrates but no products were detected by LC-MS analysis. Auxiliary redox partners, including cognate ferredoxin FdxE (*Rv1786*)– reductase (FprA or Fdr), and ferredoxin reductase– ferredoxin pairs from different sources, as well as hydrogen peroxide to bypass the requirements for molecular oxygen and NADPH-derived reducing equivalents were also tested. The design of assays to detect enzymatic activity with CYP143 is complicated by the lack of information on its natural substrate. Despite unsuccessful attempts with enzymatic activity, we set up crystallization trials with G334A mutant and RIF.

The structure of the G334A mutant in complex with RIF was determined by X-ray crystallography at 1.07 Å (Fig. 2 and Table S2). The CYP143 structure has a typical cytochrome P450 fold. Binding of RIF does not induce significant conformational changes to the overall structure of CYP143. We also determined the ligand-free structure of the wild type protein at 1.00 Å resolution. The structures of the ligand-free and RIF-bound CYP143 have an all-atom rmsd of 0.259 Å.

**Figure 2.**
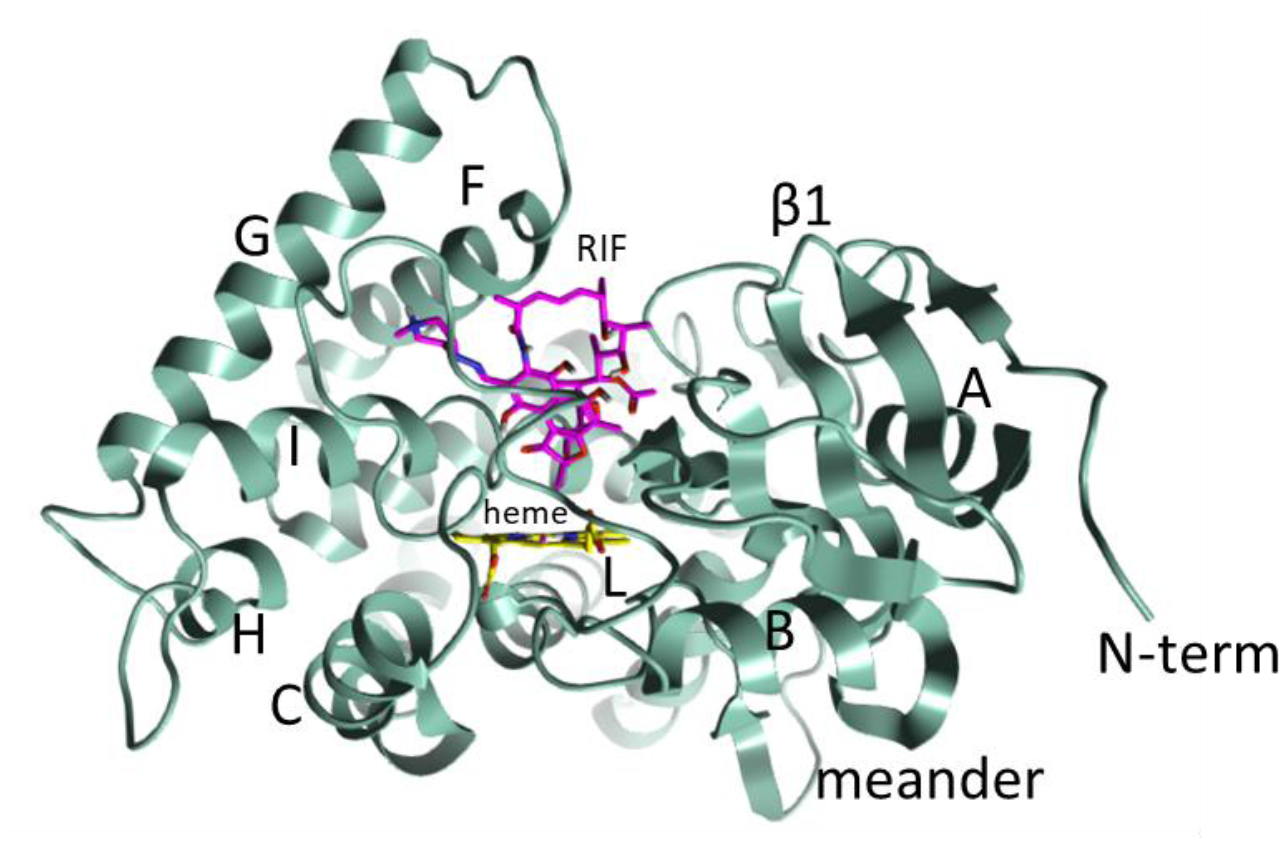
Overall structure of Gly334Ala mutant of CYP143 of *M.tuberculosis*.

The RIF binding mode explains the absence of spectral response as the ligand does not displace a water molecule from coordination with the heme iron. This water mediates the only polar interaction between the RIF and the heme. The interaction is realized via the RIF oxygen of the ansa chain between C12 and C29 of the macrocycle. The water molecule is additionally stabilized by the hydrogen bond with the main chain of highly conserved Ala232 that is a part of the oxygen-binding motif (**A**/G)GX(D/E)T in the I-helix of P450. The Thr236 of the motif is also making hydrophobic contacts with the RIF. The RIF molecule adopts a “closed”-type structure with polar groups pointing towards the naphthoquinone core, which increases the lipophilicity. The hydroxyls of the ansa chain (at C21 and C23) and the hydroxyls of the naphthoquinone (at C1 and C8) (Fig. 3 and 5) are on the same side of the molecule. The electron density is missing for the methylpiperazine branch suggesting flexibility of this part of the drug.

**Figure 3.**
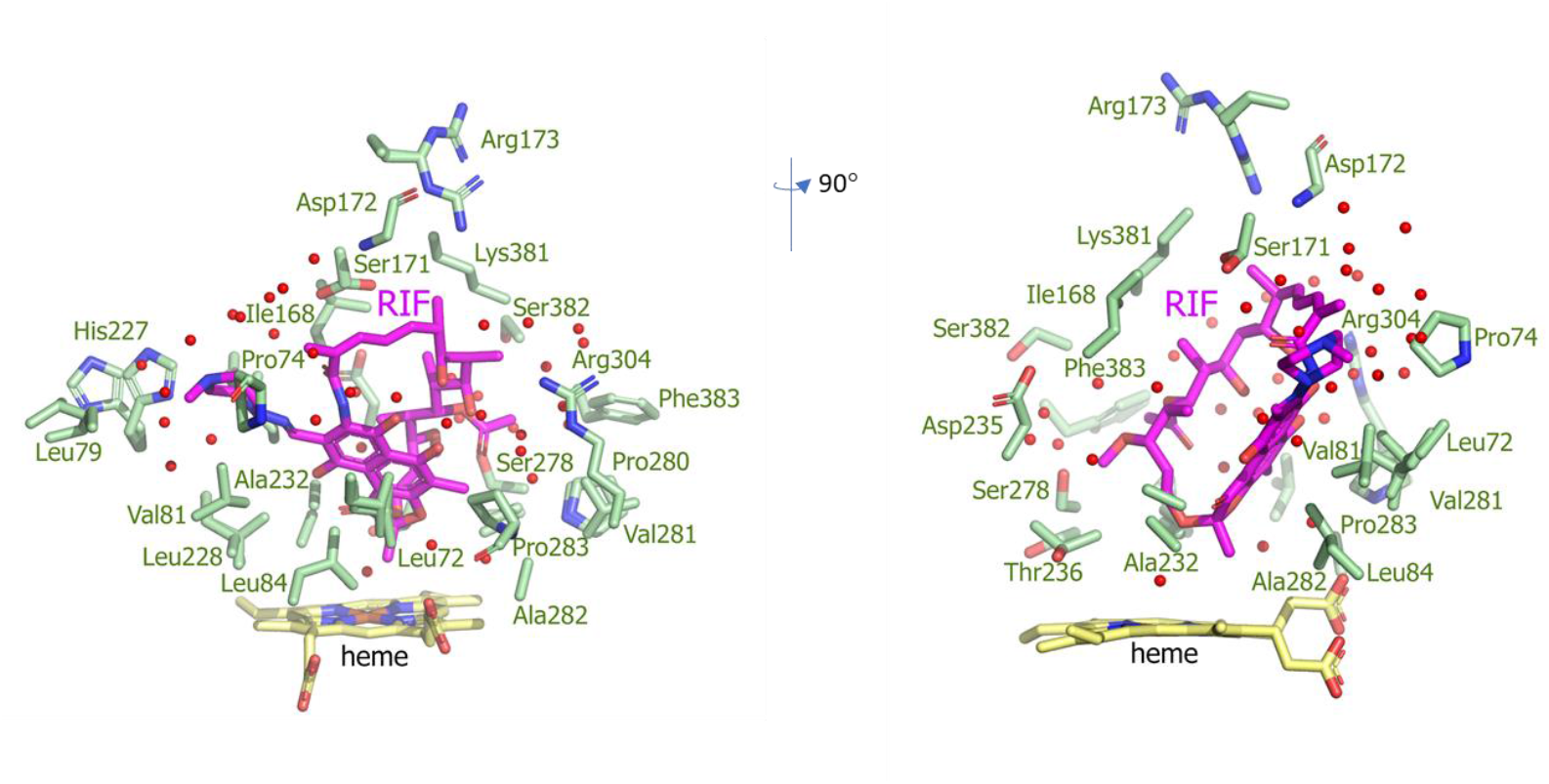
Two views of RIF binding in the active site of CYP143. Heme and RIF are shown as yellow and magenta sticks, respectively. The residues 4 Å around RIF are labeled, the water molecules are shown as red spheres.

Binding of RIF to CYP143 involves numerous van der Waals contacts with hydrophobic side chains of Leu72, Pro74, Leu79, Val81, Leu84, Ile168, Leu228, Leu231, Ala279, Pro280, Ala282, Pro283, Phe383 and the heme (Fig. 3). Proline residues from B’ helix (Pro74 and Pro75) and the loop between K helix and the β1−4 strand (Pro280 and Pro283) are imposing conformational restrictions and defining the shape of the active site cavity from these two substrate recognition sites (SRS1 and SRS5 according to^27^. The RIF naphthoquinone binds at ∼60° angle above the heme propionates, whereas the ansa chain runs almost parallel to the naphthoquinone and spans from the top of the active site cavity, formed by F-G loop and the loop near the C-terminus (SRS6), to the I helix.

A lot of water molecules remain in the active site cavity (Fig. 3), most likely they enter the active site along with RIF surrounding the ligand. Some water molecules (WAT47, WAT726 and WAT732) might be important for maintaining the “closed” conformation of RIF similarly to those described in solution ^28^,while the others are concentrated close to the surface. Overall, RIF conformation perfectly fits into the active site cavity (Fig. 4B). The closest to iron C13 atom of RIF is placed 4.6 Å away and this methyl group is pointed towards the pyrrole ring D of the heme.

**Figure 4.**
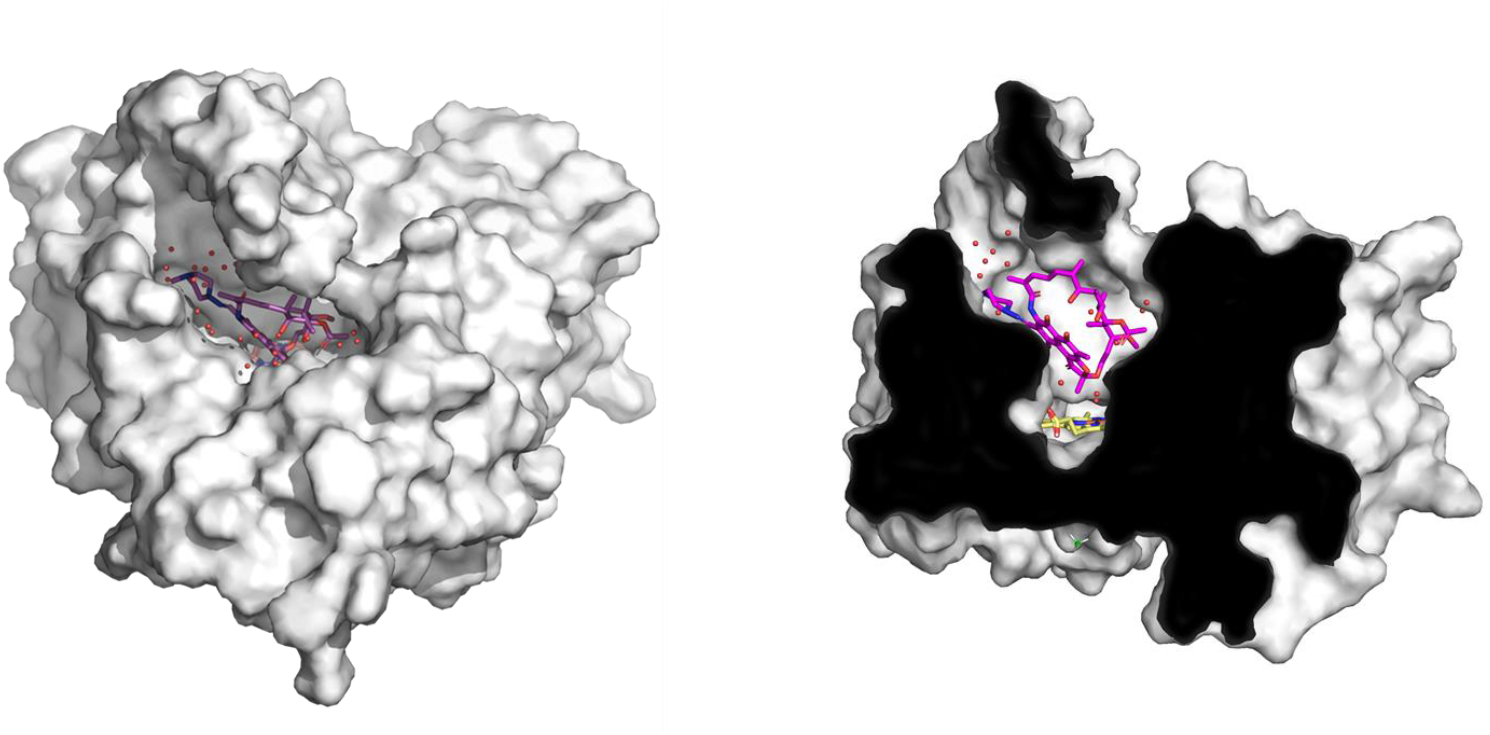
The view of RIF binding from the distal surface (left panel). The cross section of the binding site (right panel). The RIF molecule is shown as magenta sticks.

Altogether, RIF binds deep in the catalytic pocket, and in this ligand-bound state CYP143 adopts an open conformation with the RIF molecule exposed to the surface (Fig. 4).

The Gly334 residue and surrounding region are conserved among slowly growing Mycobacteria species (Fig. S3). The mutated residue G334A is located on the proximal surface of the protein where redox partner binding occurs during the catalytic cycle. The residue is below the heme and in the loop preceding the highly conserved heme-binding loop crucial for anchoring the heme and catalytic activity. To check if the additional methyl group of Ala334 residue restricts local conformational flexibility and thus affects ligand binding we analyzed its immediate surrounding. Ala334 seems to fit well into the space surrounded by residues Pro277, Val273 and His332 and does not seem to directly affect ligand binding (Fig. 5). To further understand how the mutation might be related to the resistance we solved the crystal structure of the WT protein in complex with RIF at 1.22 Å resolution. The corresponding Gly334 residue local environment only slightly changes compared with the Ala mutant. The overall structure and the binding mode of RIF are not affected as well. The mutation might be important for the catalytic activity as it is adjacent to conserved Phe335 in other P450 isoforms controlling redox potential and electron transfer to the heme^29^, but our structural data cannot clearly differentiate the role of this mutation.

**Figure 5.**
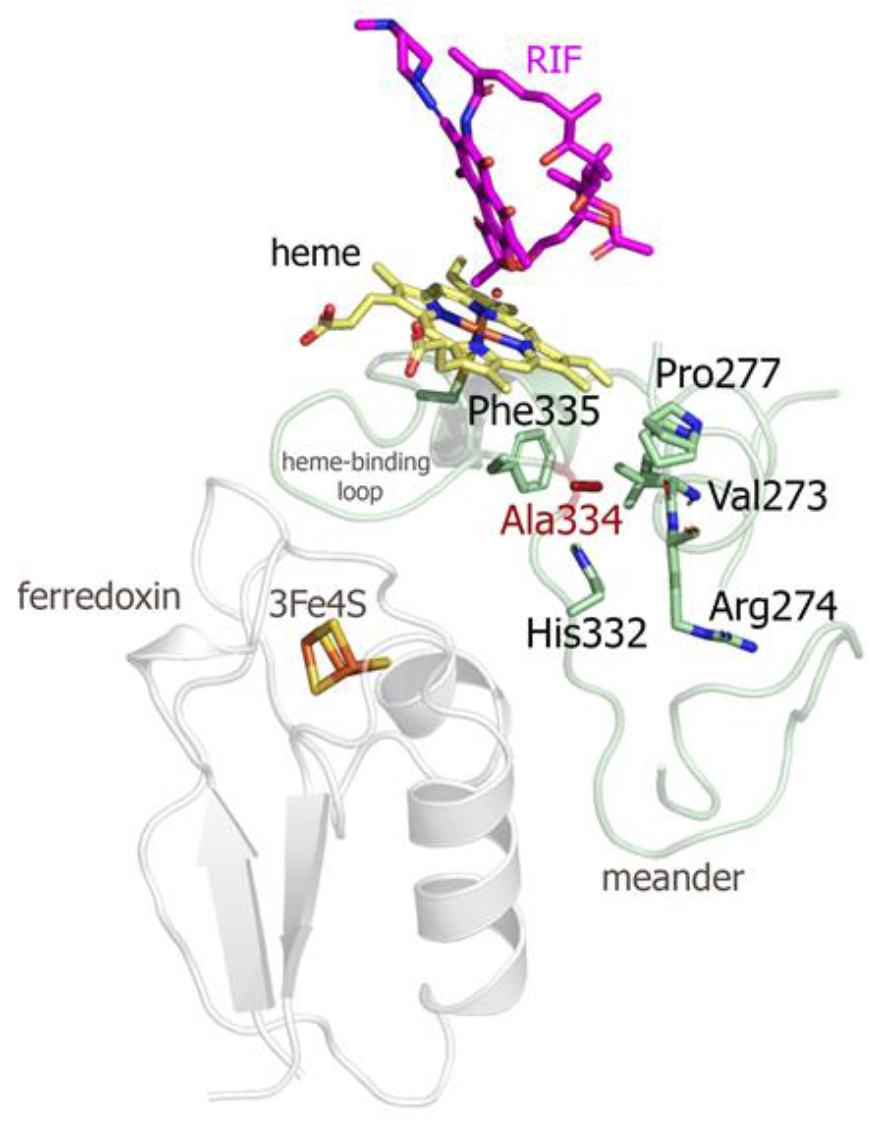
The local environment of the mutated residue Ala334. The residue is highlighted as a dark red stick. The ferredoxin is taken from the structure of the ternary complex to visualize the interaction elements on the proximal surface of CYP143.

To verify the ligand binding mode, we used Rifamycin S as a natural compound from a bacterial source^30^. Rifamycin S has a quinone that can participate in a reversible oxidation-reduction involving two electrons. In our structural studies with Rifamycin S we used both WT and G334A CYP143 proteins and solved structures at 1.13 Å and 1.12 Å resolution, respectively. Rifamycin S binds exactly the same way as RIF and similarly, the Ala334 side chain does not affect the ligand binding.

For other derivatives we used a wild type protein. The high-resolution crystal structures with other rifamycins – drug rifaximin and two synthetic derivatives – compounds 36200 and 61054 (Fig. 6) confirmed that all compounds bind at the same site above the heme via water. The electron densities for all compounds enabled unambiguous placement of ligands (Fig. S7−S9). The structural differences between used rifamycins induce only minor changes in the contacts between each compound and CYP143. It should be noted that compounds having longer radicals at C3-C4 such as rifabutin and synthetic derivatives with additional aromatic ring and/or modifications at C8 were not accommodated in the active site as we were unable to get respective complex structures.

**Figure 6.**
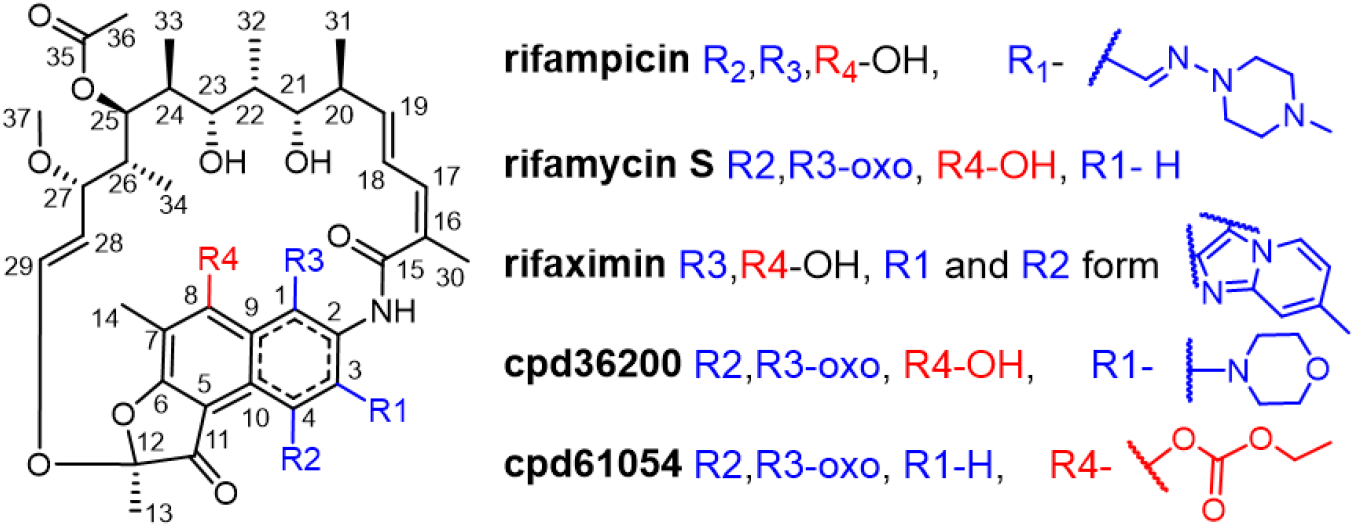
The Markush structure of rifamycins for which crystal structures in complex with CYP143 are solved in this study.

To understand if the redox partner affects the ligand binding for CYP143 we used CYP143– ferredoxin fusion protein we previously designed^18^. The ternary complex CYP143–Rv1786– 36200 was determined by X-ray crystallography to a resolution of 2.30 Å (Fig. 7). The ligand binding did not induce significant conformational changes in the protein–protein complex (PDB ID: 8AMQ), the distance between redox centers preserved, and the water-mediated interaction of the ligand with heme iron remained unchanged. In the active site compound 36200 interacts with the same residues as RIF. The overlay of the ternary structure CYP143–Rv1786–36200 with the CYP143–36200 complex reveals the same differences previously observed between the fusion protein CYP143–Rv1786 (PDB ID: 8AMQ) and CYP143 alone (PDB ID: 8AMO)^18^. In particular, the presence of the redox partner affects the positioning of the meander on the proximal surface and B’ helix flexibility (the density for the Asn77 residue is missing) on the distal side of the heme. In both ligand-free and 36200-bound fusion structures on the proximal surface His332 interacts with Glu270, while His330 interacts with Glu271 and Asp326 bringing the meander close to the K helix. The meander loop (residues Asp326-His332) adopts the conformation that accommodates ferredoxin binding.

**Figure 7.**
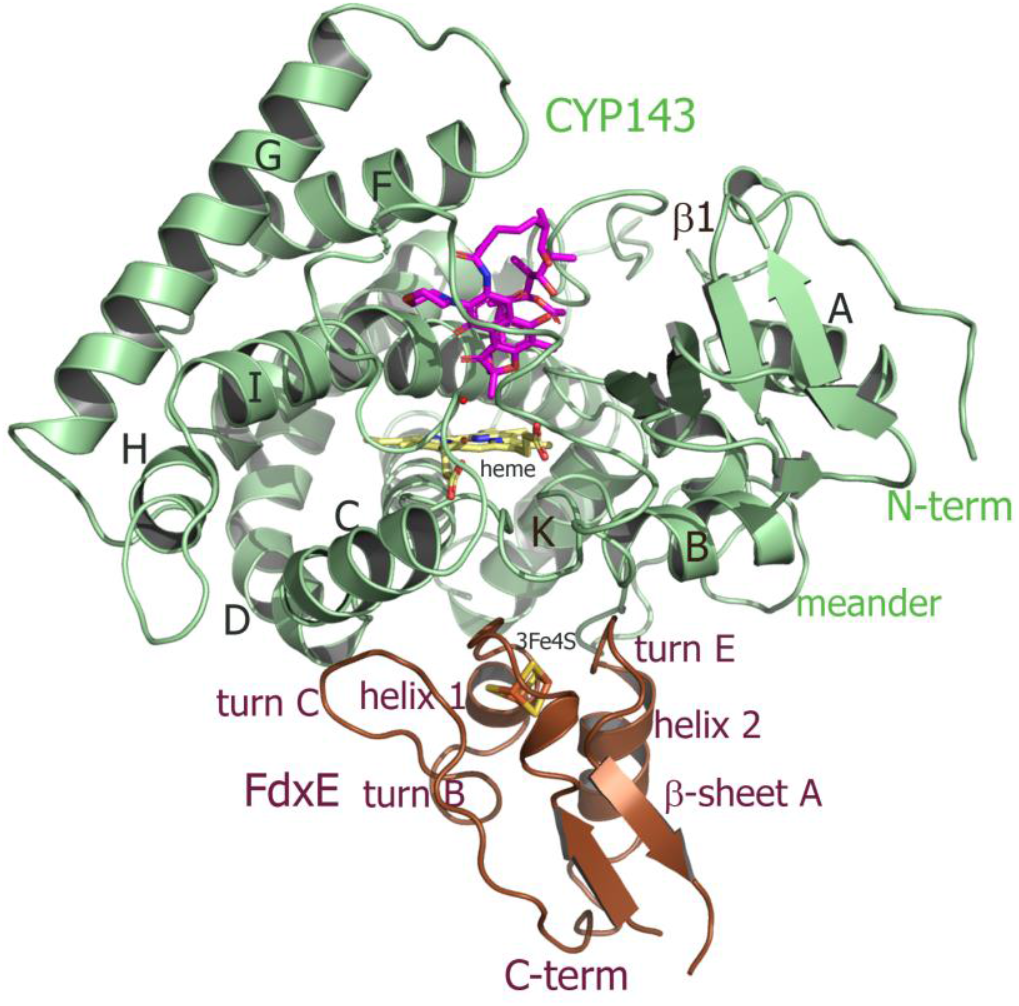
Overall structure of the FdxE–CYP143 fusion protein in complex with compound 36200. FdxE binds (brown) at the proximal face of CYP143 (green). The shortest distance between the CYP143 heme iron and the ferredoxin [3Fe–4S] cluster is 16.7 Å.

## Discussion

The key points we could deduce from the analysis of the obtained crystal structures with atomic-level information about Rif, rifamycin S and synthetic derivatives bound to CYP143 can be summarized as follows.

1. The Rif anchoring to CYP143 occurs via water molecule mediating interaction between the ligand and the iron centre of the heme. The similar mode of binding was observed in the structure of CYP121–cYY complex, where initial substrate binding does not induce the water displacement from the heme coordination^31,32^. CYP121 catalyzes C-C bond formation yielding mycocyclosin, a specific compound with yet unknown role in Mtb biology, and the cross-coupling reaction mechanism is still debated^33^. The observed binding mode should be taken into consideration for ligand screening targeting other CYP proteins using UV-Vis spectral assays.
2. The mode of ligand binding suggests a previously unseen site of metabolism/transformation. The majority of modifications of the ansa bridge (e.g. ribosylation or glucosylation of hydroxyl groups at C21 or C23, hydroxylation of C30 or C31, hydrogenation of some double bonds, epoxidation of C18–C19) have a detrimental effect on the anti-microbial activity of the derivatives^34, 35^, while modifications at C3 and C4 produce more active semisynthetic drugs. The enzyme-catalyzed modifications of rifamycin antibiotics also include N-hydroxylation by flavoproteins such as rifampicin monooxygenase (RIFMO)^36^ and linearization (i.e., degradation) by monooxygenases ROXs^37^. However, none are described for C13 and/or for the oxygen connecting a flexible ansa bridge with the aromatic core.
3. In complex with CYP143 RIF and derivatives adopt “closed” ansa bridge conformation which is distinct from an “open” ansa bridge conformer of lower lipophilicity crucial for binding with the main target - bacterial RNA polymerases ^38, 39, 40, 41^. This conformation has higher lipophilicity and enables diffusion in bacterial cell membranes^28^. Indeed, the CYP143 active site is hydrophobic and complementary in shape to tested rifamycins. Notably, the selective binding of RIF in “closed” conformation was not captured in RIF-inactivation enzymes. For example, RIF phosphotransferase (PDB ID: 5HV1), different rifampicin monooxygenases (PDB ID: 5KOX; 6BRD), rifampin ADP-ribosyl transferase (PDB ID: 2HW2), all have “open” conformation of RIF. The ligand conformational selection is well known for CYP proteins ^42,43^ and most likely this “closed” ligand conformation in CYP143 is specific for a CYP-mediated metabolism.
4. The ligands in this conformation can easily enter the open active site of CYP143 and reach the heme cofactor not affecting protein conformation. Typically, CYP proteins adopt closed conformation upon ligand binding. The preserved open conformation of CYP143 even in the presence of the redox partner ferredoxin might be a feature to accommodate large ligands. Alternatively, temporal or permanent sequestration of rifampicin/s in the CYP143 active site might be a mechanism to decrease active concentration of the drug and ultimately lead to resistance.

*M. tuberculosis* enters the body via the lower respiratory tract, where it interacts with alveolar macrophages and microbiota. It should be noted that the lung microbiome is quite distinct from the gut microbiome as its major role is resisting infection and regulating immunity^44^. The upper and lower respiratory tracts differ in their microbial composition and biomass. Moreover, the lower airway microbiome is frequently changing, balancing between microbial immigration and microbial clearance^13^. The core lung microbiome includes Pseudomonas, Streptococcus, Prevotella, Fusobacterium, Haemophilus, Veillonella, and Porphyromonas^45,13,46^. Recent data obtained using systematic culture and whole genome sequencing of the airway commensal bacteria from healthy subjects^44^ revealed that clinically relevant genes that encode antimicrobial synthesis are present in many genera. For example, rifB (rifamycin polyketide synthase) gene present in 20 isolates (*Veillonella and Staphylococcus spp*.). Although the presence of this gene does not necessarily indicate the rifamycin synthesis, this potential might be important for understanding the Mtb interaction with the local microbiome. We speculate that during invasion Mtb might also utilize enzymes involved in antibiotic binding/degradation, such as CYP143, to outcompete the microbiome of the lower respiratory tract. This hypothesis warrants further investigation.

## Materials and methods

*Bioinformatics*. The methodology for building CYPome phylogenetic trees was taken from elsewhere ^47^. Briefly, the sequence of CYP143 of *Mtb* H37Rv was used as a query to build a set of CYPs by BLASTP and TBLASTN searches against the GenBank database. The accession numbers are provided in Figure S1. The resulting sequence sets with added Rv1786 sequence were aligned with MEGA software (version 12.0.8) ^48^ using MUSCLE. The sequences were analyzed using Bayesian inference implemented on BEAST v2.7.7 ^49^. The JTT model with a discrete gamma (G) distribution across sites (four categories) was selected with MEGA software (version 12.0.8) ^48^ as a good fitting one for phylogenetic analysis of the P450s. The uncorrelated relaxed log normal clock model and a Yule birth model with birth rate gamma distribution (α = 0.001, β = 1000) were chosen to allow every branch of a tree to evolve with its own rate. The analysis was performed by running 10^7^ generations in two chains, keeping every 1000th tree. The burn-in percentage 20 was used to sum the results of the runs, and the consensus tree was created in TreeAnnotator. The final tree was edited in FigTree v.1.4.4, and only *cyp* genes were depicted ^50^, available from http://tree.bio.ed.ac.uk/software/figtree/.

The number of CYPs in different species and presence of specific CYPs in the genomes were either described with UniProt (https://www.uniprot.org/), NCBI GenBank (https://www.ncbi.nlm.nih.gov/), TB (http://tbdb.bu.edu/tbdb_sysbio/MultiHome.html) databases or Mycobrowser (https://mycobrowser.epfl.ch/), and checked with article ^51^. Certain CYP isoforms could not be fully presented here as we rely on available annotations at the moment and they might not be fulfilled yet. The presence of CYP143 and Rv1786 was also checked or specified with the Bacterial and Viral Bioinformatics Resource Center (BV-BRC, https://www.bv-brc.org/). To study the evolutionary history of the *Mycobacterium* genus we used a methodology described in a similar study ^47^. In short, 16S rRNA sequence of *Mtb* H37Rv was used as a query to obtain the 16S rRNAs of 32 bacterial species by a BLASTN search against the RefSeq database. The sequence sets were aligned with MEGA software (version 12.0.8) ^48^ using MUSCLE. The phylogenetic analysis was performed using BEAST v2.7.7 ^49^. The alignment was analyzed with 10^7^ generations of MCMC under the HKY substitution model and with gamma distributed rates (5 categories), uncorrelated relaxed log normal clock model, and a Yule birth model with birth rate gamma distribution (α = 0.001, β = 1000). The burn-in percentage 20 was used to sum the results of the runs, and the consensus tree was formed in TreeAnnotator and edited in FigTree v.1.4.4 ^50^, available from http://tree.bio.ed.ac.uk/software/figtree/.

*Site-directed mutagenesis* was performed using Toyobo KOD-plus mutagenesis kit according to the manufacturer’s protocol. The resulting plasmid harboring G334A mutant gene was verified by DNA sequencing.

*Expression and purification of CYP143 WT and G334A mutant* was performed as previously described ^18^.

### Spectral binding studies

To determine azole binding constants (apparent Kd values) optical titration was performed using dual-beam spectrophotometer Jasco V-760 (Jasco Inc, Japan) in 1-cm path length quartz cuvettes. Compound stock solutions 10mM were made in DMSO. 1-2μM of CYP143 was used per titration in 50 mM potassium phosphate buffer, pH 7.5 at room temperature. Titration was repeated at least three times and Kd was calculated as described previously ^47^.

### HTS ligand binding assay

The interaction of CYP143 with the ligands was monitored by the spectral response in a 50 mМ potassium phosphate buffer, pH 7.4. in the 350–500 nm wavelength range using multi-mode microplate reader CLARIOstar Plus (BMG LABTECH). The final protein concentration was 3 μM. The final compound concentration was 100 μM (added from a 10 mM stock solution in DMSO). The assay was performed in the 96-well UV transparent plate at RT.

### Rifampicin–CYPs binding analysis

Rifampicin (final concentration 60 μM) was added to the protein solution at different concentrations (15, 30, 45, and 60 μM) of CYP143. The final samples were mixed gently, incubated for 5 min and centrifuged for 20 min using centrifugal filters with a 30 kDa molecular weight cut-off. UV–visible absorption spectra of the filtrates were recorded on a Cary 5000 UV–Vis–NIR spectrophotometer (Agilent Technologies, USA) in 1-cm path length quartz cuvettes at 300–600 nm wavelength range. The same procedure was applied to a rifampicin solution alone as a reference to assess rifampicin filtration through the membrane. CYP141 and CYP51 from *Mycobacterium tuberculosis* were used as negative controls. All experiments were carried out at room temperature in 50 mM Tris–HCl buffer, pH 7.4 containing 300 mM NaCl. The stock solution of rifampicin was prepared in the same buffer. The concentration of rifampicin was determined using an extinction coefficient of ε_475_ = 15.279 M^−1^ · cm^−1^.

### Crystallization of CYP143 and FdxE–CYP143

CYP143 (150 µM) and FdxE–CYP143 (200–250 µM) were crystallized in 96-well plate using a sitting-drop method with commercially available kits from Qiagen (NeXtal Classics II screen) and Molecular Dimensions (Structure screens 1 and 2) at 20 °C with 1:1 protein/mother liquor ratio with the ligand concentration of 1-2 mM. The synthetic derivatives of rifampicin were purchased from InterBioScreen ltd., catalog ID: STOCK1N-61054 and STOCK1N-36200. Red colored crystals of CYP143 appeared overnight and FdxE−CYP143 in 2 weeks. The best crystals of CYP143 and FdxE−CYP143 grew in 0.2 M Sodium chloride, 0.1 M Bis-Tris pH 6.0, 25% (w/v) PEG3350.

### Data collection, and X-ray structure determination

The obtained crystals were harvested, and diffraction data were collected at various synchrotron beamlines (see Table S1), with all data sets acquired at 100 K under cryogenic conditions. Diffraction images were processed and scaled using the XDS software suite^52^. The data quality and potential anomalies were assessed using phenix.xtriage^53^.

Phases were obtained by molecular replacement in PHENIX Phaser^54^, employing initial polyalanine models derived from previously determined structures (PDB: 8AMO and 8AMQ)^18^, which served as references for the CYP143 and chimeric CYP143-Rv1786 datasets, respectively. Each dataset contained one molecule per asymmetric unit. Model refinement was performed through iterative cycles combining phenix.refine^55^ and manual model building in Coot^56^. Restraints for non-standard ligands (rifampicin derivatives) were generated using the Grade webserver and incorporated into subsequent refinement rounds. High-quality electron density maps enabled unambiguous ligand placement without further modification.

Final models were validated using phenix.molprobity^57^ and subjected to Quality Control Check v3.2. Each ligand derivative displays an overall real-space correlation coefficient (RSCC) >0.90 indicating a reliable fit to the electron density (Fig. S7-S9). Full data collection and refinement statistics are summarized in Table S2.

## Author contributions

Conceptualization − A.G. and N.S.; Methodology − K.B., V.B., A.G., N.S.;

Investigation − A.G., C.M, T.V., M.K., I.G., V.K., T.S., A.K., V.B., N.S.;

Writing – Original Draft. A.G and N.S.;

Writing – Review & Editing. A.G., V.B., N.S.;

Funding Acquisition − A.G., V.B, N.S.;

Supervision − A.G., K.B., V.B., N.S.

## Data availability

Models of ligand free CYP143 WT and G334A mutant, CYP143 WT and G334A in complex with rifampicin, CYP143 WT and G334A in complex with rifamycin S, CYP143 WT in complex with rifaximin and compound 36200, fusion protein CYP143–Rv1786–36200 are being deposited to PDB.

## Acknowledgements

We thank Dr. Kurt Wollenberg, Dr. Andrei Gabrielian and Dr. Michael Allen Harris for their help with data retrieval from the TB portal. We also thank Dr. Leonid Kaluzhskiy and Dr. Evgeniy Yablokov for their efforts with the SPR analysis, Dr. Sergey Osipenko and Dr. Yury Kostyukevich for LC-MS analysis. We thank LNLS, ESRF, PETRAIII and SSRF for providing beamtime for X-ray data collection.

## Funding and additional information

Synchrotron data collection and data treatment were supported by the Russian Ministry of Science and Higher Education (agreement #075-15-2025-512). VB and ChM acknowledge the Ministry of Science and Higher Education of the Russian Federation (agreement # 075-03-2025-662, project FSMG-2024-0012) for support of structural analysis. This research was funded by the Belarusian Republican Foundation for Fundamental Research, grant number X23RNF-090. The work was performed in part within the framework of the Program for Basic Research in the Russian Federation for a long-term period (2021-2030, №122030100168-2).

