## Supplemental Information for "True-atomic-resolution insights into the interactions of antibiotic rifampicin and rifamycin derivatives with orphan CYP143 of *M.tuberculosis*"

**Supplementary Information.**

**Figure S1.** Detailed phylogenetic tree of P450 protein sequences for *Mycobacteria*. Colors represent different groups of CYP records. The NCBI reference sequence accession numbers are provided for all used records.

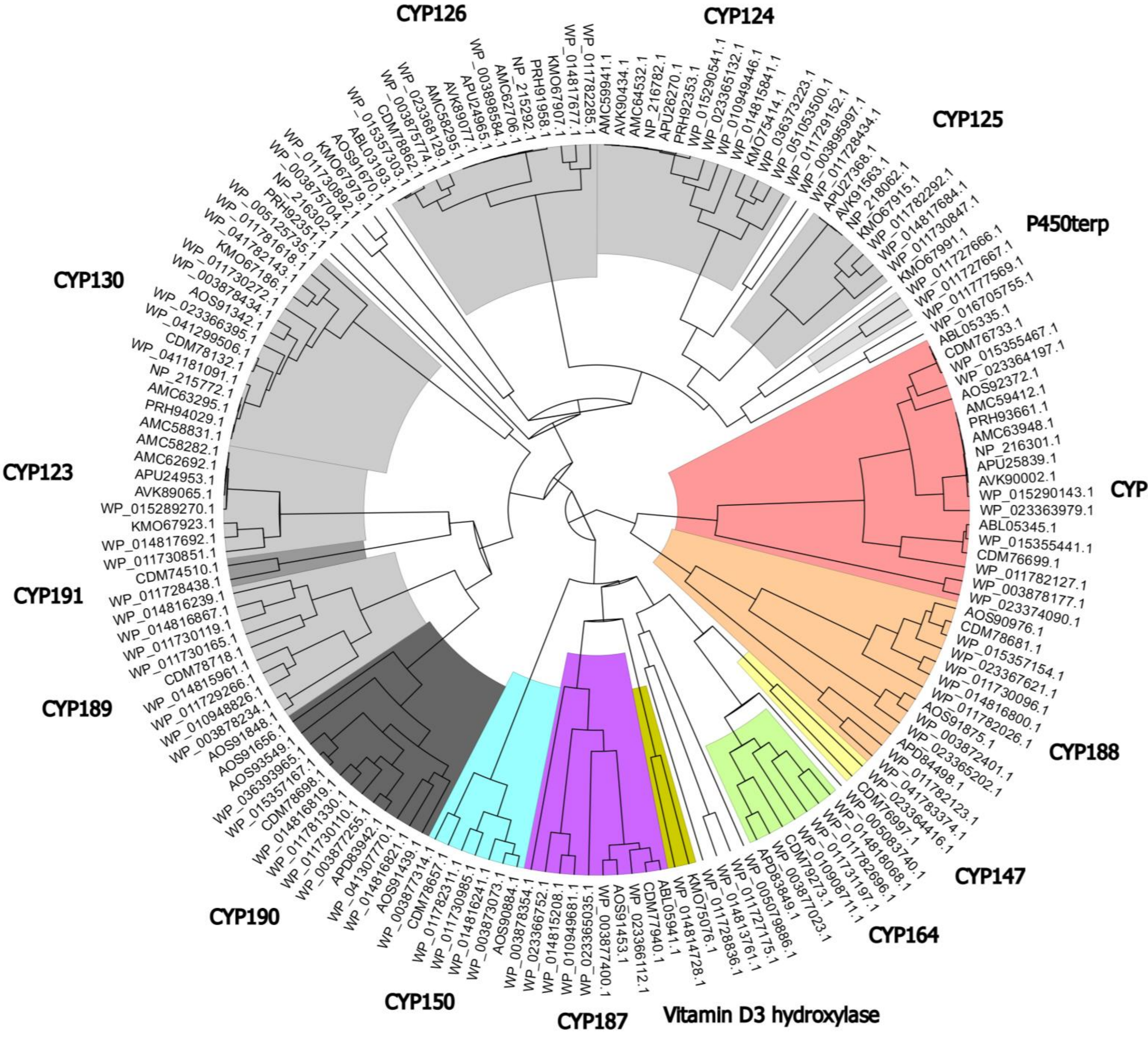

**Figure S2.** Synteny of CYP143 (Rv1785c) for selected *Mycobacteriaceae* taxonomy genomes. The SynTax web server[1] was used with default parameters, and CYP143 protein query (NP_216301.1) was used as an input and shown in bold. The organisms are sorted by decreasing TBLASTN scores. Both paralogs and orthologs are color coded, when present. The genes of *esx5* export system are also highlighted in blue, while Rv1786 (Fdx, ferredoxin) and Rv1785c (CYP143) are in pink.

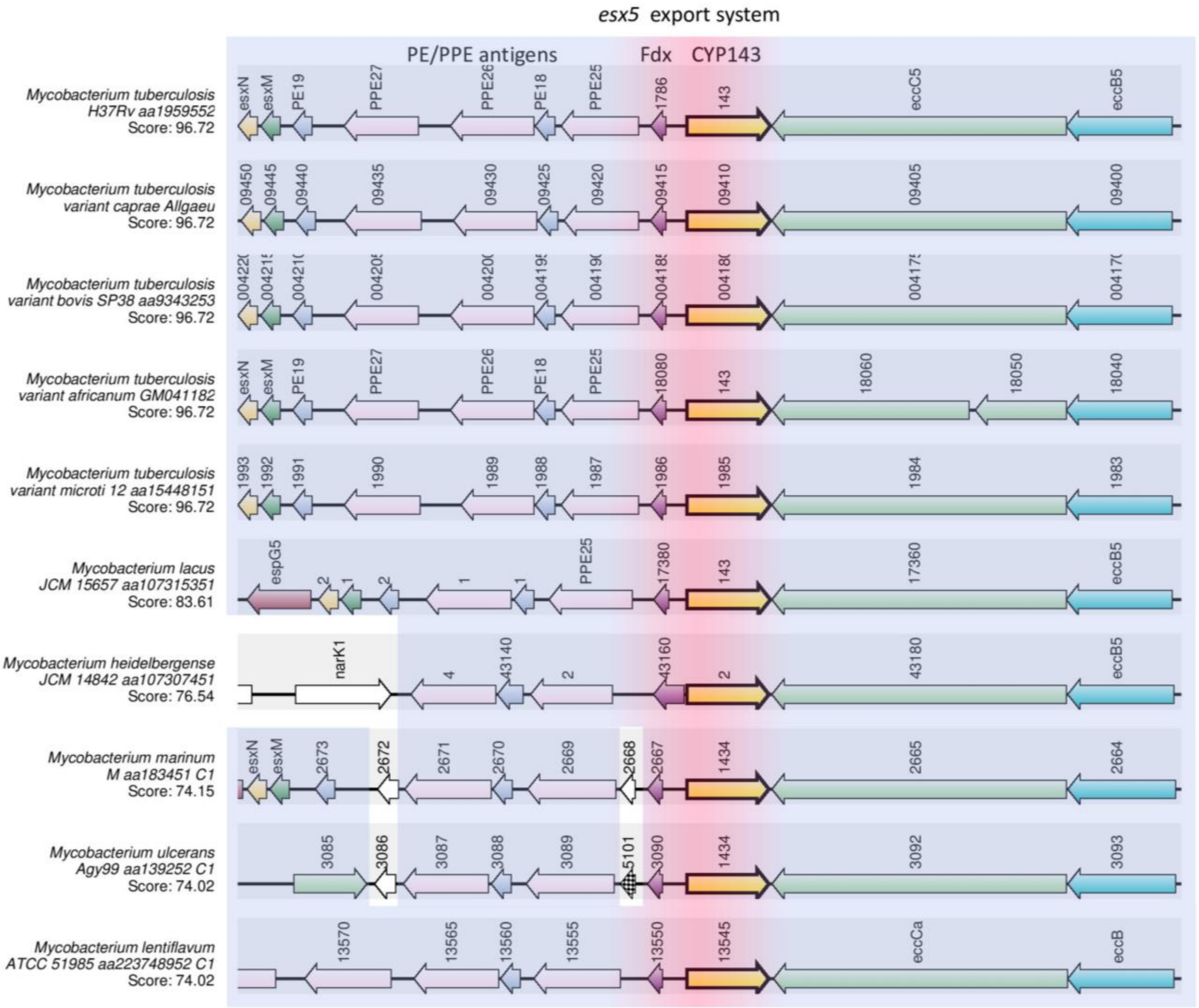

**Figure S3.** Multiple sequence alignment of CYP143 obtained from BLASTP and genomic context search of CYP143 in slowly growing *Mycobacteria*. Blue color gradient indicates percentage identity, where deep blue defining a high percentage. Records from BLASTP and SyntTax[1] search were merged in one .fasta file, which was aligned with ProbconsWS algorithm in Jalview software (version 2.11.5.0)[2]. The NCBI reference sequence accession numbers are provided below for all used records.

NP_216301.1 cytochrome P450 Cyp143 [Mycobacterium tuberculosis H37Rv]
CCC26877.1 putative cytochrome P450 143 CYP143 [Mycobacterium tuberculosis variant africanum GM041182]
AMC59412.1 cytochrome P450 Cyp143 [Mycobacterium tuberculosis variant microti]
APU25839.1 cytochrome [Mycobacterium tuberculosis variant caprae]
QEF47374.1 cytochrome P450 [Mycobacterium tuberculosis variant bovis]
WP_023363979.1 cytochrome P450 [Mycobacterium kansasii]
WP_085158134.1 cytochrome P450 [Mycobacterium lacus]
WP_083077239.1 cytochrome P450 [Mycobacterium heidelbergense]
WP_012394383.1 cytochrome P450 [Mycobacterium marinum]
WP_011740947.1 cytochrome P450 [Mycobacterium ulcerans]
WP_085225328.1 cytochrome P450 [Mycobacterium florentinum]
WP_042911914.1 cytochrome P450 [Mycobacterium intracellulare]
WP_076101432.1 cytochrome P450 [Mycobacterium colombiense]
WP_085256727.1 cytochrome P450 [Mycobacterium saskatchewanense]
WP_067167332.1 cytochrome P450 [Mycobacterium marseillense]
WP_163708547.1 cytochrome P450 [Mycobacterium timonense]
WP_083065961.1 cytochrome P450 [Mycobacterium arosiense]
WP_083099034.1 cytochrome P450 [Mycobacterium mantenii]
WP_104183227.1 cytochrome P450 [Mycobacterium avium]
WP_085239037.1 cytochrome P450 [Mycobacterium europaeum]
WP_085268789.1 cytochrome P450 [Mycobacterium parmense]
WP_015355441.1 cytochrome P450 [Mycobacterium liflandii]

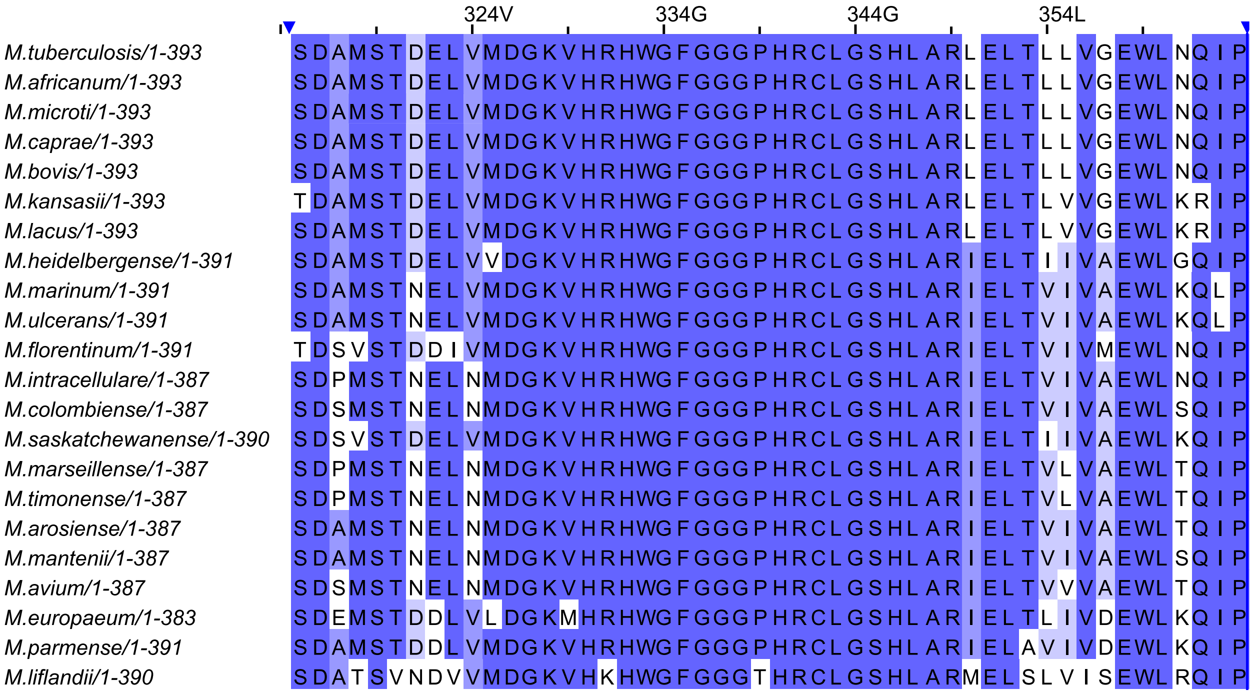

**Figure S4.** ITC binding curve of CYP143 with rifamycin S in 50 mM potassium phosphate buffer, pH 7.4, 5% DMSO. Rifamycin S binding to CYP143 was enthalpy-driven and weak. No binding was detected for rifampicin or rifabutin. The low solubility of rifampicin, rifabutin, and rifamycin S in aqueous solutions posed challenges for ensuring sufficient concentration during ITC experiments.

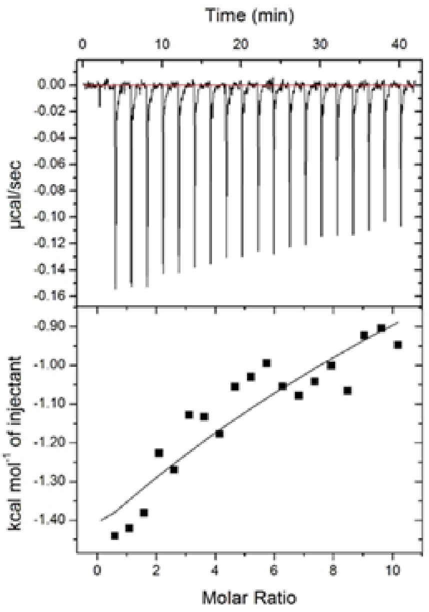


Experiments were performed on an ITC200 calorimeter equipped with the data acquisition and analysis software ORIGIN 7 (MicroCal Inc.) at 25ºC. To study the interactions between rifampicin, rifabutin, and rifamycin S and CYP143, protein was dialyzed overnight against 50 mM potassium-phosphate buffer, pH 7.4, containing 5% DMSO. An aliquot of 20 µM CYP143 was placed in the calorimetric cell and titrated with 500 µM rifampicin or rifamycin S or rifabutin dissolved in the same buffer. The first injection (1 µl, omitted from the analysis) was followed by 20 injections of 4 µl at 2-minute time intervals. The titration syringe was continuously stirred at 750 rpm, and the temperature of the calorimetric cell was 25 °C. Injection of the rifamycin S into the buffer alone was used as reference titration. The data were analyzed with the program ORIGIN (MicroCal) using a one-site binding model.

**Figure S5.** ITC binding curves of CYP143 with azoles in 50 mM potassium phosphate buffer, pH 7.4, 5% DMSO. A) clotrimazole, B) fluconazole, C) econazole, D) miconazole, E) ketoconazole, F) imazalil.

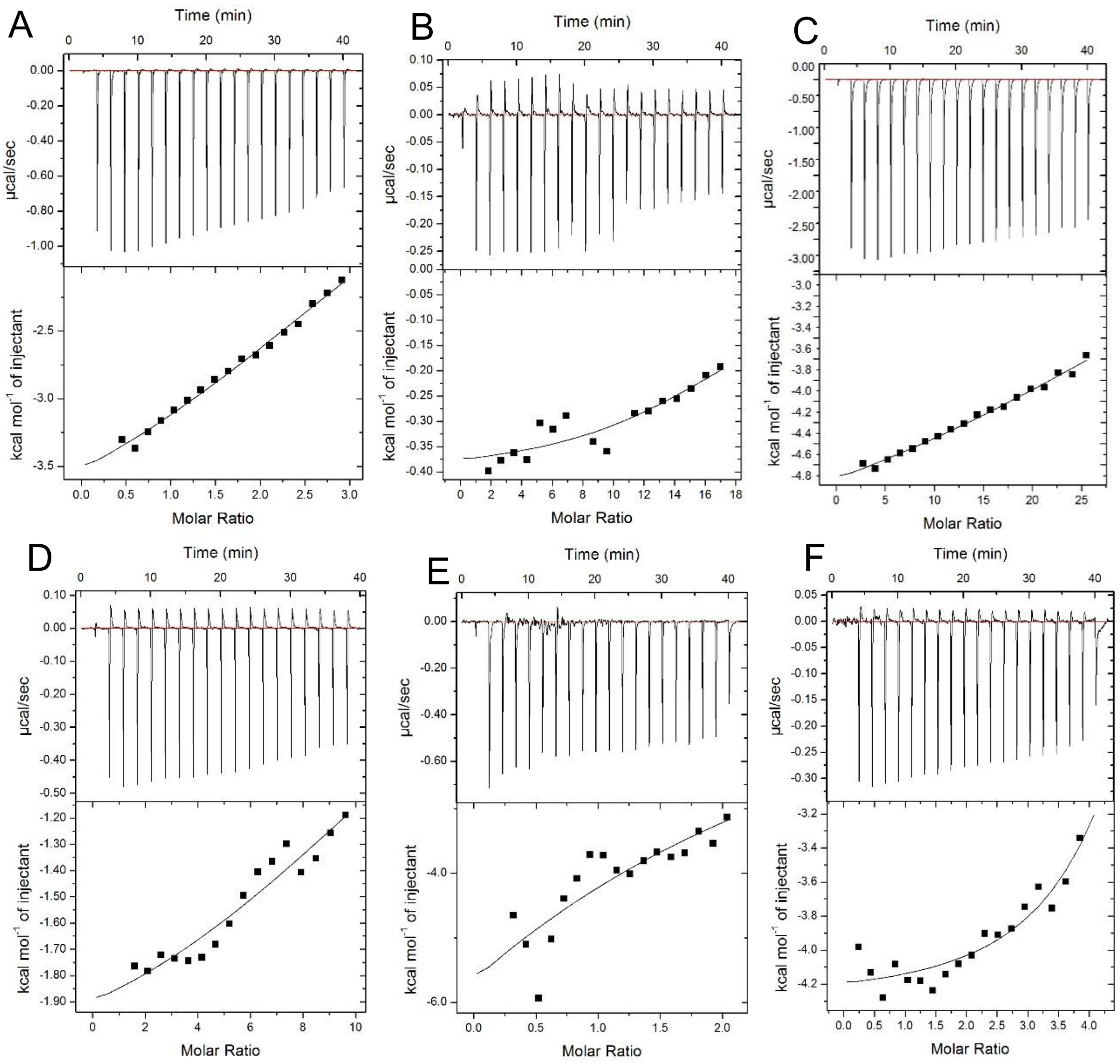


**Table S1.** Thermodynamic parameters of the interaction between CYP143 and azoles

| **CYP143** | **Kd, µmol** | **ΔH, kcal/mol** | **-TΔS (kcal/mol)** | **ΔG, kcal/ mol** |
|---|---|---|---|---|
| Clotrimazole | 83.6±6.7 | -3.9 | -1.1 | -5.0 |
| Fluconazole | 23.6±3.8 | -2.4 | -1.8 | -4.2 |
| Econazole | 110±8.1 | -4.4 | -2.2 | -6.6 |
| Miconazole | 28.9±6.1 | -4.2 | -3.6 | -7.8 |
| Ketoconazole | 54.0±3.1 | -3.3 | -3.1 | -6.4 |
| Imazalil | 3.4±0.6 | -12.2 | 3.4 | -8.8 |

Experiments were performed on an ITC200 calorimeter equipped with the data acquisition and analysis software ORIGIN 7 (MicroCal Inc.) at 25ºC. To study the interactions between azoles and CYP143, protein was dialyzed overnight against 50 mM potassium-phosphate buffer, pH 7.4, containing 5% DMSO. An aliquot of 20-50 μM CYP143 was placed in the calorimetric cell and titrated with 500-2500 μM azoles dissolved in the same buffer. The first injection (1 μl, omitted from the analysis) was followed by 20 injections of 4 μl at 2-minute time intervals. The titration syringe was continuously stirred at 750 rpm, and the temperature of the calorimetric cell was 25 °C. The data were analyzed with the program ORIGIN (MicroCal) using a one-site binding model.

**Table S2.** Summary of data collection and refinement statistics. Statistics for high-resolution shells are given in parentheses.

| PARAMETERS | CYP143 ligand free (MT) | CYP143 ligand free (WT) | CYP143-RifamycinS (WT) | CYP143-Rifamycin S(MT) | CYP143-Rifampicin (WT) | CYP143-Rifampicin (MT) | CYP143-Rifaximin (WT) | CYP143-36200 (WT) | CYP143Rv1786-3620 (WT) | CYP143-61054 (WT) |
|---|---|---|---|---|---|---|---|---|---|---|
| Beamline | LNLS (MANACA) | ESRF (ID23EH1) | ESRF (ID23-1) | LNLS (MANACA) | PETRAIII (P13) | PETRAIII (P13) | ESRF (ID23-1) | SSRF (BL17UM) | SSRF (BL17UM) | SSRF (BL17UM) |
| Wavelength, Å | 0.9772 | 0.9720 | 0.7749 | 0.9772 | 1.0332 | 0.9762 | 0.7749 | 0.9792 | 0.9792 | 0.9792 |
| Resolution range, Å | 30.74 - 1.10 (1.13 - 1.10) | 47.57-1.00 (1.03-1.00) | 37.43-1.13 (1.16-1.13) | 46.39 - 1.12 (1.15 - 1.12) | 47.83 - 1.22 (1.25 - 1.22) | 47.46 - 1.07 (1.10 - 1.07) | 37.56 - 1.21 (1.24 - 1.21) | 47.33 - 1.56 (1.60 - 1.56) | 32.93 - 2.31 (2.42 - 2.31) | 30.62 - 1.54 (1.58 - 1.54) |
| Space group | P 1 | P 1 | P 1 | P 1 | P 1 21 1 | P 1 | P1 | P 1 | C 1 2 1 | P 1 |
| Unit-cell, Å | 42.31 49.19 54.40 112.94 98.47 109.31 | 42.26 48.98 54.3 112.11 99.2 109.32 | 42.08 48.60 54.05 113.23 99.53 109.71 | 42.8 53.16 54.4 61.54 67.82 73.8 | 52.97 75.43 54.19 90 118.04 90 | 42.17 48.88 54.21 112.64 98.59 109.15 | 42.13 48.57 53.89 111.39 99.29 109.58 | 42.04 47.98 53.77 111.46 99.2 109.89 | 92.9 55.31 98.92 90 92.81 90 | 42.09 49 54.14 112.46 98.37 109.7 |
| Total reflections | 470715 (26800) | 1196534 (74763) | 457165 (34047) | 486075 (32244) | 747686 (54782) | 519122 (32556) | 361770 (21009) | 166553 (9550) | 150170 (22364) | 180934 (13258) |
| Unique reflections | 134805 (8462) | 187389 (13144) | 127472 (9252) | 138099 (9046) | 106894 (7696) | 142844 (9357) | 102517 (6594) | 46682 (2942) | 22195 (3218) | 50167 (3651) |
| Multiplicity | 3.5 (3.2) | 6.4 (5.7) | 3.6 (3.7) | 3.5 (3.6) | 7.0 (7.1) | 3.6 (3.5) | 3.5 (3.2) | 3.6 (3.2) | 6.8 (6.9) | 3.6 (3.6) |
| Completeness, % | 91.6 (78.0) | 96.30 (91.39) | 95.68 (92.36) | 92.61 (80.49) | 95.74 (92.75) | 90.35 (79.60) | 94.5(81.9) | 94.28 (80.01) | 99.81 (99.82) | 94.76 (92.53) |
| Mean $I/\sigma(I)$ | 20.57 (6.42) | 18.33 (1.24) | 7.27 (0.6) | 12.79 (1.14) | 15.55 (1.27) | 15.38 (1.15) | 9.13 (0.50) | 9.74 (2.11) | 7.25 (0.82) | 7.08 (0.63) |
| $CC_{1/2}$, % | 99.9 (96.7) | 100(66) | 99.8 (26.3) | 100 (53.8) | 99.9 (73.4) | 100 (62.3) | 99.9 (23.4) | 99.6 (66.7) | 99.7 (52.9) | 99.7 (34.7) |
| Reflections used in refinement | 134794 (3713) | 187370 (5858) | 127277 (4124) | 138073 (3981) | 106845 (3493) | 142803 (3820) | 101952 (2480) | 46671 (2227) | 22180 (2739) | 50117 (2725) |
| Reflections used for $R_{free}$ | 6813 | 9416 | 6478 | 6766 | 5217 | 7222 | 5208 | 2360 | 1115 | 2548 |
| $R_{work}/R_{free}$ | 0.1040/0.1263 | 0.1041/0.1212 | 0.1296/0.1692 | 0.1365/0.1614 | 0.1359/0.1741 | 0.1119/0.1429 | 0.1441/0.1796 | 0.1283/0.1675 | 0.2088/0.2455 | 0.1665/0.1999 |
| Number of. non-H atoms | 4426 | 4748 | 4460 | 4186 | 4584 | 4739 | 4604 | 4332 | 3737 | 4011 |
| Macromolecules | 3514 | 3725 | 3399 | 3262 | 3591 | 3685 | 3518 | 3333 | 3501 | 3397 |
| Ligands | 48 | 44 | 108 | 93 | 107 | 106 | 103 | 109 | 118 | 126 |
| Solvent | 864 | 979 | 953 | 831 | 886 | 948 | 983 | 890 | 118 | 488 |
| Protein residues | 384 | 384 | 385 | 385 | 389 | 385 | 385 | 385 | 448 | 385 |
| RMS(bonds) | 0.007 | 0.007 | 0.003 | 0.006 | 0.008 | 0.012 | 0.013 | 0.006 | 0.046 | 0.005 |
| RMS(angles) | 0.97 | 0.89 | 0.72 | 0.89 | 1.07 | 1.12 | 1.02 | 0.86 | 1.72 | 0.81 |
| Ramachandran favored (%) | 97.91 | 97.91 | 98.16 | 98.17 | 98.45 | 97.39 | 98.68 | 98.16 | 97.51 | 97.89 |
| Ramachandran allowed (%) | 2.09 | 2.09 | 1.84 | 1.83 | 1.55 | 2.61 | 1.32 | 1.84 | 2.49 | 2.11 |
| Ramachandran outliers (%) | 0.00 | 0.00 | 0.00 | 0.00 | 0.00 | 0.00 | 0.00 | 0.00 | 0.00 | 0.00 |
| Rotamer outliers (%) | 0.75 | 0.74 | 0.27 | 0.84 | 0.50 | 0.49 | 0.26 | 0.55 | 0.54 | 0.54 |
| Clashscore | 2.08 | 3.94 | 2.53 | 3.52 | 1.86 | 3.11 | 2.05 | 2.73 | 3.63 | 2.53 |
| Average B-factor | 17.44 | 22.54 | 23.33 | 22.40 | 31.27 | 21.96 | 26.16 | 21.03 | 61.60 | 31.75 |
| Macromolecules | 14.53 | 19.08 | 18.74 | 18.61 | 26.36 | 17.43 | 22.24 | 17.45 | 62.01 | 30.75 |
| Ligands | 7.88 | 9.75 | 18.52 | 15.48 | 30.84 | 23.04 | 18.67 | 16.45 | 53.49 | 36.13 |
| Solvent | 29.81 | 36.26 | 40.25 | 38.04 | 51.24 | 39.42 | 40.90 | 35.02 | 57.63 | 37.78 |

**Figure S6.** 2Fo-Fc maps (1.5 $\sigma$) of the Rifampicin-bound CYP143. A and B refer to the wild type and mutant CYP143, respectively. The ligand is colored according to the B factor.

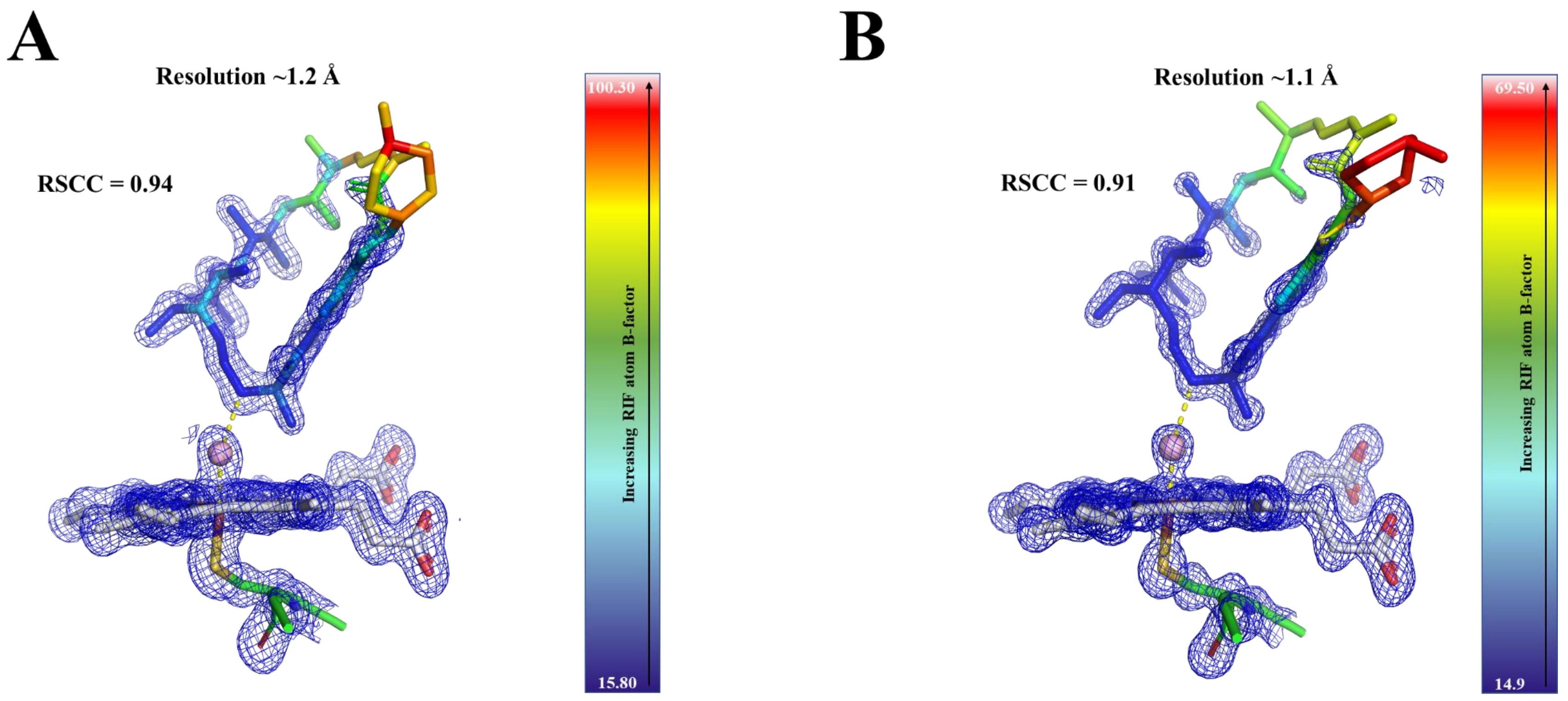

**Figure S7.** 2Fo-Fc maps (1.5 $\sigma$) of the Rifamycin S-bound CYP143. A and B refer to the wild type and mutant CYP143, respectively. The ligand is colored according to the B factor.

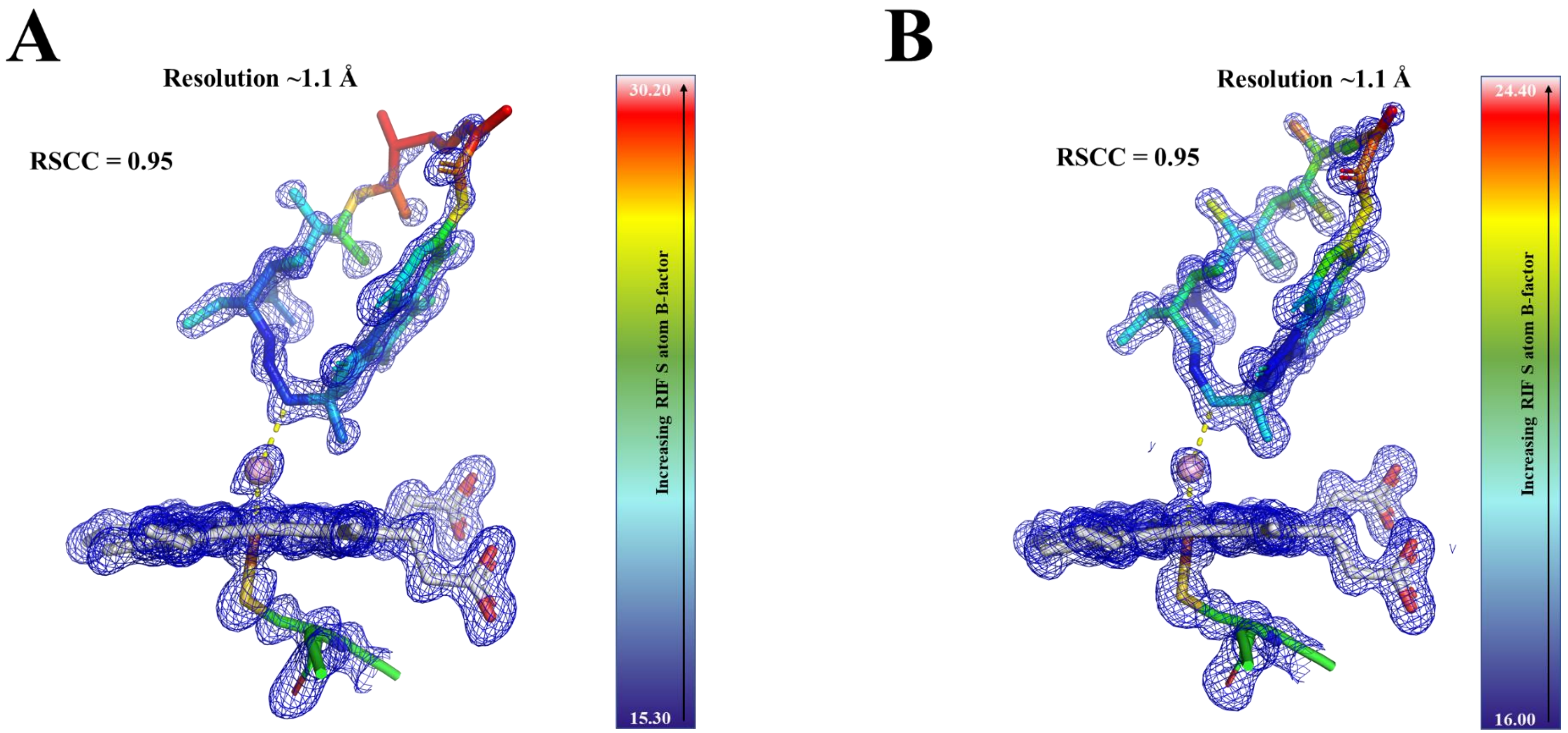

**Figure S8.** 2Fo-Fc maps (1.5 $\sigma$) of Rifaximin (A) and 61054 (B) interacting with the CYP143. The ligands are colored according to the B factor.

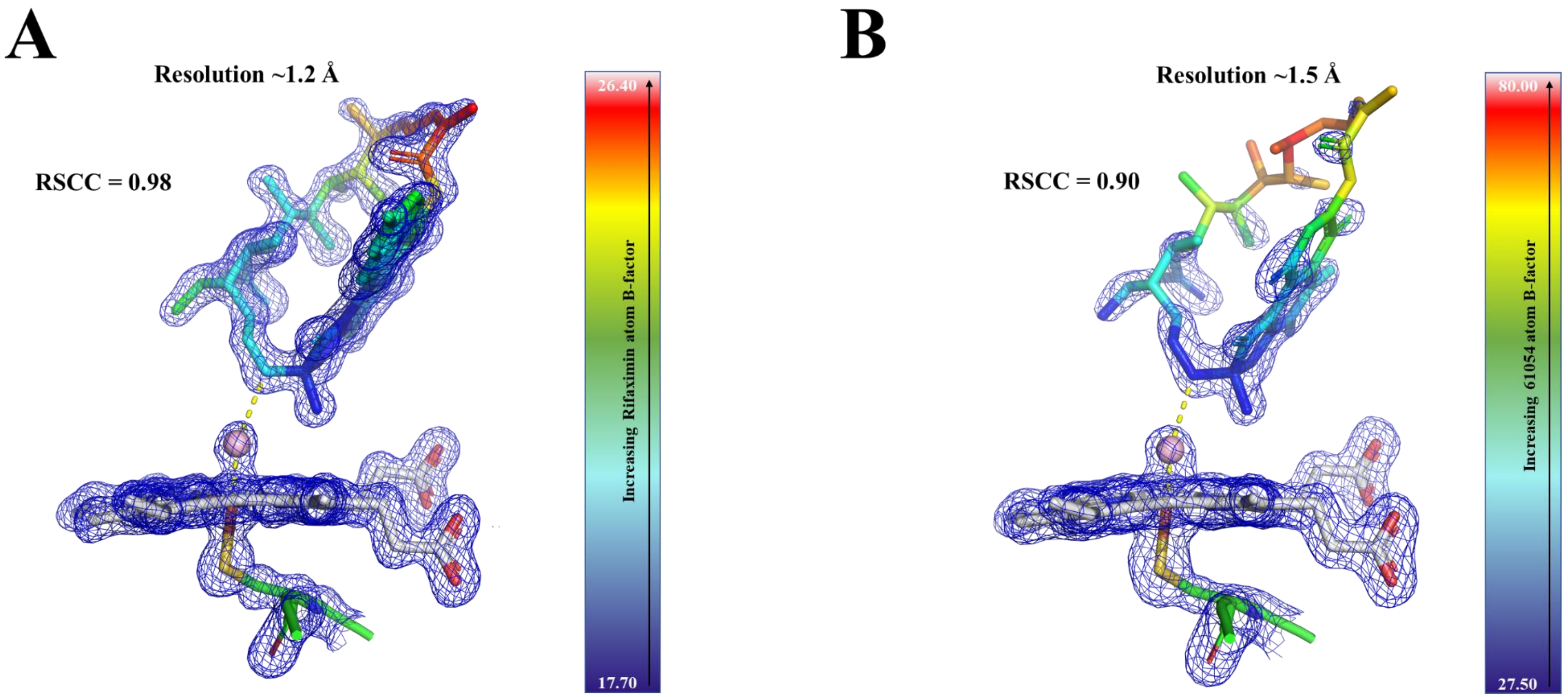

**Figure S9.** 2Fo-Fc maps (1.5 $\sigma$) of the compound 36200 interacting with CYP143 (A) and CYP143-FdxE complex (B). The ligand is colored according to the B factor.

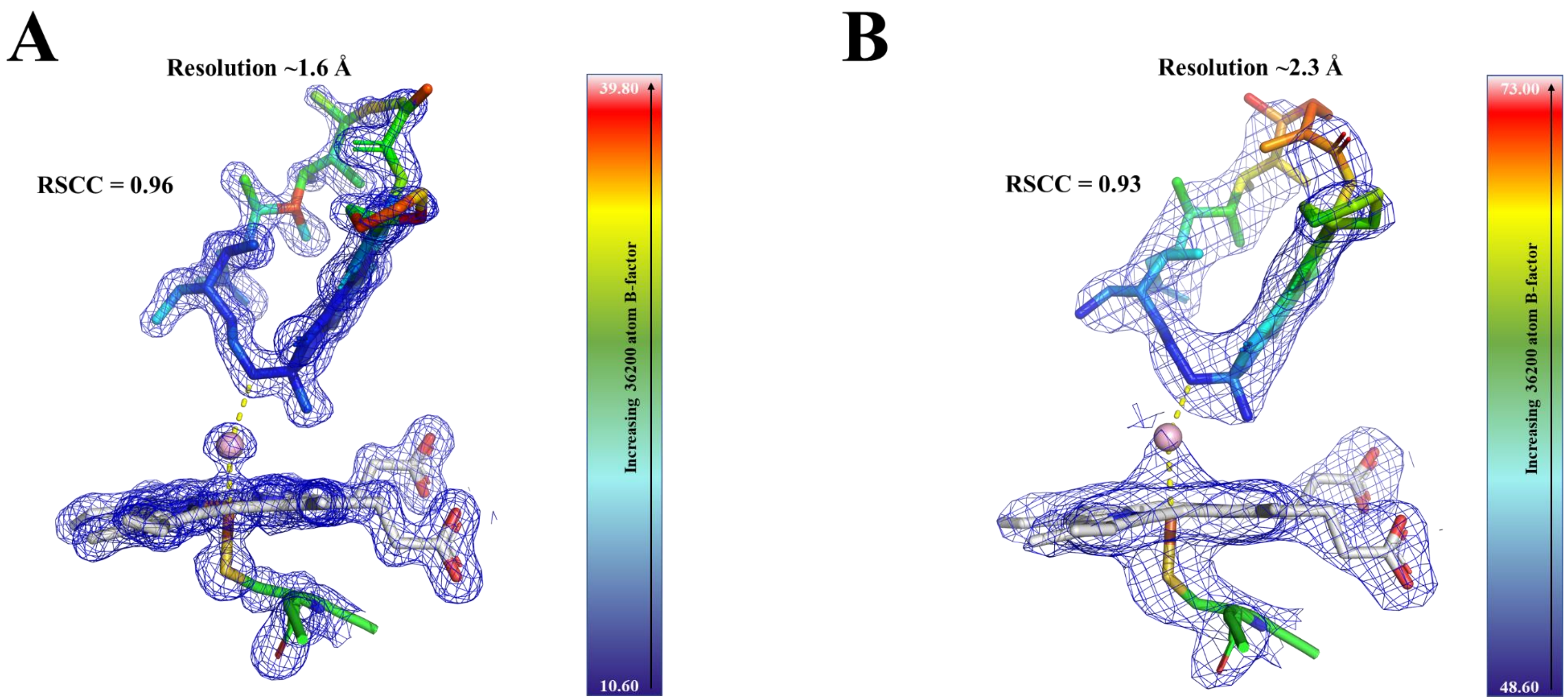
